# Mutational analysis of LtgC, a lytic transglycosylase required for cell separation in *Neisseria gonorrhoeae*

**DOI:** 10.1101/2023.06.20.545760

**Authors:** Ryan E. Schaub, Krizia Perez-Medina, Joshua Tomberg, Robert A. Nicholas, Joseph P. Dillard

**Author notes:** Address correspondence to: Joseph P. Dillard, 1550 Linden Drive, Madison, WI 53706.

## Abstract

Lytic transglycosylases function to degrade peptidoglycan strands that comprise the bacterial cell wall. Degradation of peptidoglycan at the septum following cell division is necessary for cell separation, and a deletion of *ltgC* in *Neisseria gonorrhoeae* results in growth in clusters of around 6-20 cells rather than as normal diplococci or monococci. *N. gonorrhoeae* LtgC is a homolog of *Escherichia coli* MltA, and comparison of the two proteins shows that LtgC has an extra domain not found in MltA, referred to as domain 3. To better understand the function of LtgC, we characterized *N. gonorrhoeae* mutants with substitutions in amino acids predicted to be necessary for enzymatic activity or amino acids predicted to be on the surface of domain 3, and we characterized a mutant lacking domain 3. All the mutants showed defects in cell separation, and the bacteria failed to release peptidoglycan-derived disaccharides into the medium. Purified LtgC proteins with the amino acid substitutions had reduced peptidoglycan degradation activity. LtgC was found to bind AmiC in bacterial 2-hybrid assays, and domain 3 mutations reduced binding. In human blood, an *ltgC* mutant showed decreased survival, suggesting the cell wall defects in the mutant make the bacteria more sensitive to innate immune system components.

**Importance:** *Neisseria gonorrhoeae* uses a smaller set of proteins for peptidoglycan breakdown compared to *Escherichia coli* or other model systems. The peptidoglycan breakdown that occurs at the septum following cell division in *N. gonorrhoeae* requires three proteins, amidase AmiC, amidase activator NlpD, and lytic transglycosylase LtgC. LtgC has an unusual structure that includes a third domain not found in related proteins. Using mutants that lacked LtgC activity or had amino acid changes in the third domain, we found that the extra domain is involved in interaction of LtgC with AmiC and that it is required for LtgC function for cell separation. All of the *ltgC* mutants examined showed reduced survival in blood, indicating the importance of LtgC activity for infection.

## Introduction

*Neisseria gonorrhoeae* is the causative agent of the sexually-transmitted disease gonorrhea and can cause severe complications in women, including pelvic inflammatory disease, chronic pelvic pain, and tubal-factor infertility [1]. These disease manifestations are due to the inflammatory response to bacterial products, and in particular, immune recognition of gonococcal peptidoglycan fragments leads to inflammatory cytokine production and ciliated cell sloughing [2, 3]. *N. gonorrhoeae* has a somewhat defective variant of AmpG, the permease for recycling of glycan-containing peptidoglycan fragments. As a result of this defect, gonococci release significant amounts of peptidoglycan fragments into the infection milieu [4].

The core peptidoglycan structure in *N. gonorrhoeae* is typical of Gram-negative bacteria. Peptidoglycan is made of alternating residues of *N*-acetylglucosamine (GlcNAc) and *N*-acetylmuramic acid (MurNAc), with a peptide attached to MurNAc having the sequence L-Ala-D-isoGlu-*m*DAP-D-Ala-D-Ala. The peptides can be 2, 3, 4, or 5 amino acids long, but in *N. gonorrhoeae* approximately 80% of the peptides are 4 amino acids long, and nearly 20% are 3 amino acids long [5, 6]. Differences between *N. gonorrhoeae* peptidoglycan and *E. coli* peptidoglycan are the presence of acetyl groups on the C6 hydroxyl on about 40% of muramic acid residues in gonococci, and the absence of any covalently attached proteins to gonococcal peptidoglycan [7, 8].

LtgC has been studied in *N. gonorrhoeae* and in the closely related species *N. meningitidis.* Cloud and Dillard found that deletion of *ltgC* resulted in a cell separation defect, increased autolysis, and no release of GlcNAc-anhydro-MurNAc disaccharides [9]. A study in *N. meningitidis* found that deletion of *ltgC* (also known as *gna33* or *mltA*) resulted in a cell separation defect, increased release of outer-membrane proteins into the medium, and an inability to cause infection in an infant rat model of meningococcemia [10]. In addition to LtgC, the other proteins found to be required for cell separation in *N. gonorrhoeae* or *N. meningitidis* are the *N*-acetylmuramyl-L-alanine amidase AmiC, its activator NlpD, and the FtsN-related protein Tpc [11–13]. Mutation of *nlpD* was shown to result in decreased survival of *N. meningitidis* in human blood [14].

Biochemical studies of the meningococcal enzyme found that it is a lipoprotein with lytic transglycosylase activity, capable of digesting sacculi to produce soluble peptidoglycan monomers and also capable of digesting poly(MurNAc-GlcNAc) into GlcNAc-anhydro-MurNAc disaccharides [15]. Crystal structures of *N. gonorrhoeae* LtgC identified residues in the active site including two aspartic acids (D405 and D393) predicted to be important for catalysis [16]. It was also shown that while *E. coli* MltA and gonococcal LtgC shared two similar domains, a 66 amino acid region of LtgC forms a third domain (domain 3) not found in the *E. coli* enzyme [16].

In this study, we examined the impacts on LtgC function of mutating the two aspartic acids predicted to be within the active site, mutating 5 amino acids present on the surface of domain 3, and deletion of domain 3 altogether. We examined enzymatic activity, peptidoglycan composition, peptidoglycan fragment release, cell separation, and protein-protein interactions. Our data reveal that both aspartic acid residues are indeed necessary for activity, and that domain 3 is involved in binding to AmiC. All of these mutants exhibited cell separation defects, altered peptidoglycan release, and reduced survival in whole blood.

## Results

### Mutation of *ltgC* results in cell separation defects

The crystal structure of LtgC showed that it is very similar in structure to *E. coli* MltA but has an extra domain not found in MltA, called domain 3 (Fig. 1) [16]. The authors speculated that domain 3 might function in gonococci to facilitate interactions with another protein. To better understand the function of LtgC and domain 3, we made one mutant in which the domain 3 encoding region was deleted (ΔDm3) and a second mutant where five residues on the surface of domain 3 were changed to alanine (5mut). The changes in the 5mut strain are K179A, L181A, R183A, L227A, and P228A (Fig. 1). In addition, we made two active site mutations, D405A and D393A, which are highly conserved in lytic transglycosylases and are predicted to be critical for enzymatic function.

**Figure 1:**
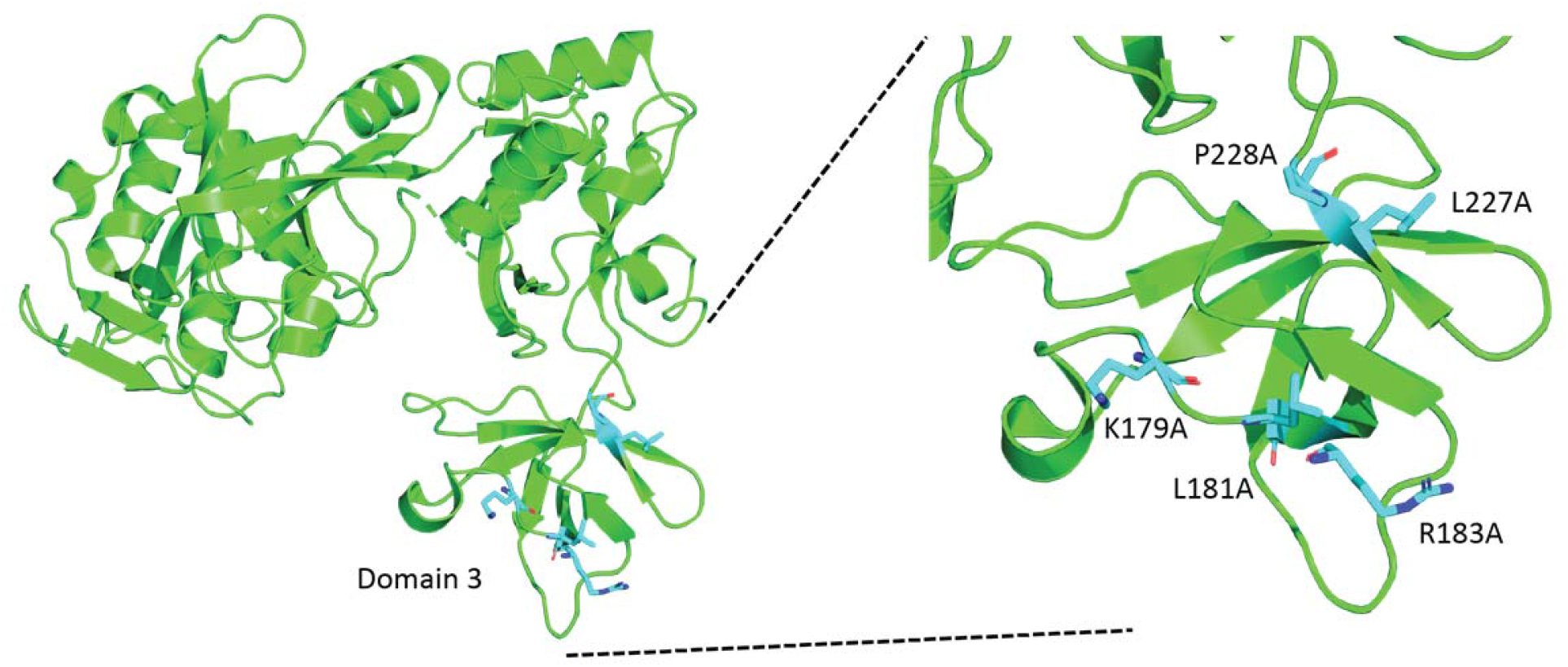
LtgC Crystal Structure. The structure of LtgC indicating the location of Domain 3 and the mutations made to create LtgC-5mut.

To examine the effects of the *ltgC* mutations, each mutation was introduced into *N. gonorrhoeae* strain FA19. Each protein also carried a C-terminal HA-tag to allow monitoring of LtgC protein levels. We first subjected the mutant strains to thin-section electron microscopy to examine cell separation. Both WT *N. gonorrhoeae* strain FA19 and a strain carrying LtgC-HA showed typical monococcal and diplococcal morphology (Fig 2A-1D). Very few aggregates of more than three cells were seen, and very few dead cells were noted. By contrast, mutants D405A or D393A showed very few monococci and diplococci, and a substantial number of lysed cells could be seen as empty cells with little to no electron-dense material inside. Most of the mutant cells were found in clusters of six or more cells, unseparated (Fig 2E-1H).

**Figure 2:**
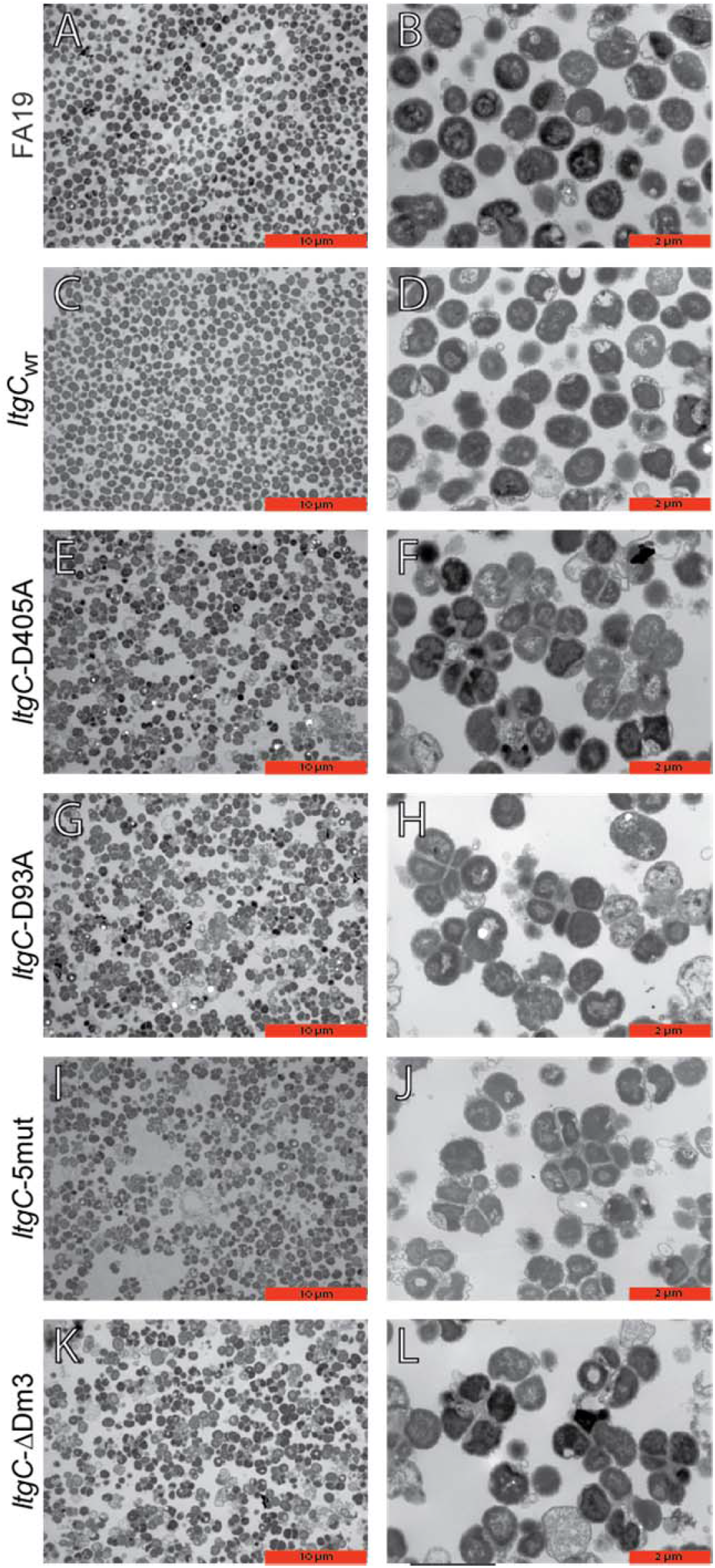
Thin-section transmission electron microscopy of *ltgC* mutants. Electron micrographs at 8,800× (A, C, E, G, H, I, and K) and 15,000× (B, D, F, H, J, and L), show that wild-type FA19 and isogenic strain with a C-terminal HA-tag (A-D) as single cocci or diplococcic. The *ltgC* mutants also contain a C-terminal HA-tag and are deficient in cell separation (E-L).

Strains harboring an *ltgC* allele in which domain 3 was deleted (*ltgC*-ΔDm3) or with 5 mutations in domain 3 (*ltgC*-5mut) showed similar phenotypes to strains with D405A or D393A active site mutations. The bacteria were observed in clusters of 6 or more cells, and lysed cells were common in each field (Fig 2I-1L).

### Mutations in *ltgC* result in altered growth characteristics

The appearance of lysed cells in the electron micrographs prompted us to examine the growth characteristics of the *ltgC* mutants. The bacteria were grown in liquid culture starting at an OD_540_=0.2, which for gonococci equates to approximately 10^8^ CFU/mL for WT strains. The WT strain carrying LtgC-HA grew to an OD_540_ of over 0.6 in six hours. By contrast, each of the *ltgC* mutants reached about OD_540_=0.3 in the same period (Fig 3A). CFU/mL values are not an accurate measurement for the unseparated bacteria growing in clusters, but the values are somewhat informative, nevertheless. The WT strain increased in CFU/mL as expected. However, the 5mut strain remained relatively unchanged in CFU numbers over the growth period, while the other mutants showed decreases in viable counts (Fig 3B). Quantification of protein in the cell pellet showed the WT strain increasing in protein amount by 2.5-fold over the growth period. The mutants showed a trend toward slower growth and reached a 2-fold increase (Fig 3C). However, it should be noted that the protein values are not statistically different for any of the mutants compared to WT. Overall, these data show that the mutants do grow in liquid culture as illustrated by the protein values, but they only accumulate a minor amount of additional clusters of cells as indicated by the OD measurements. The decrease in CFU/mL for the mutants suggests that the growth occurs by adding cells to the clusters and some of the clusters undergoing increased lysis as observed in the electron microscopy studies.

**Figure 3:**
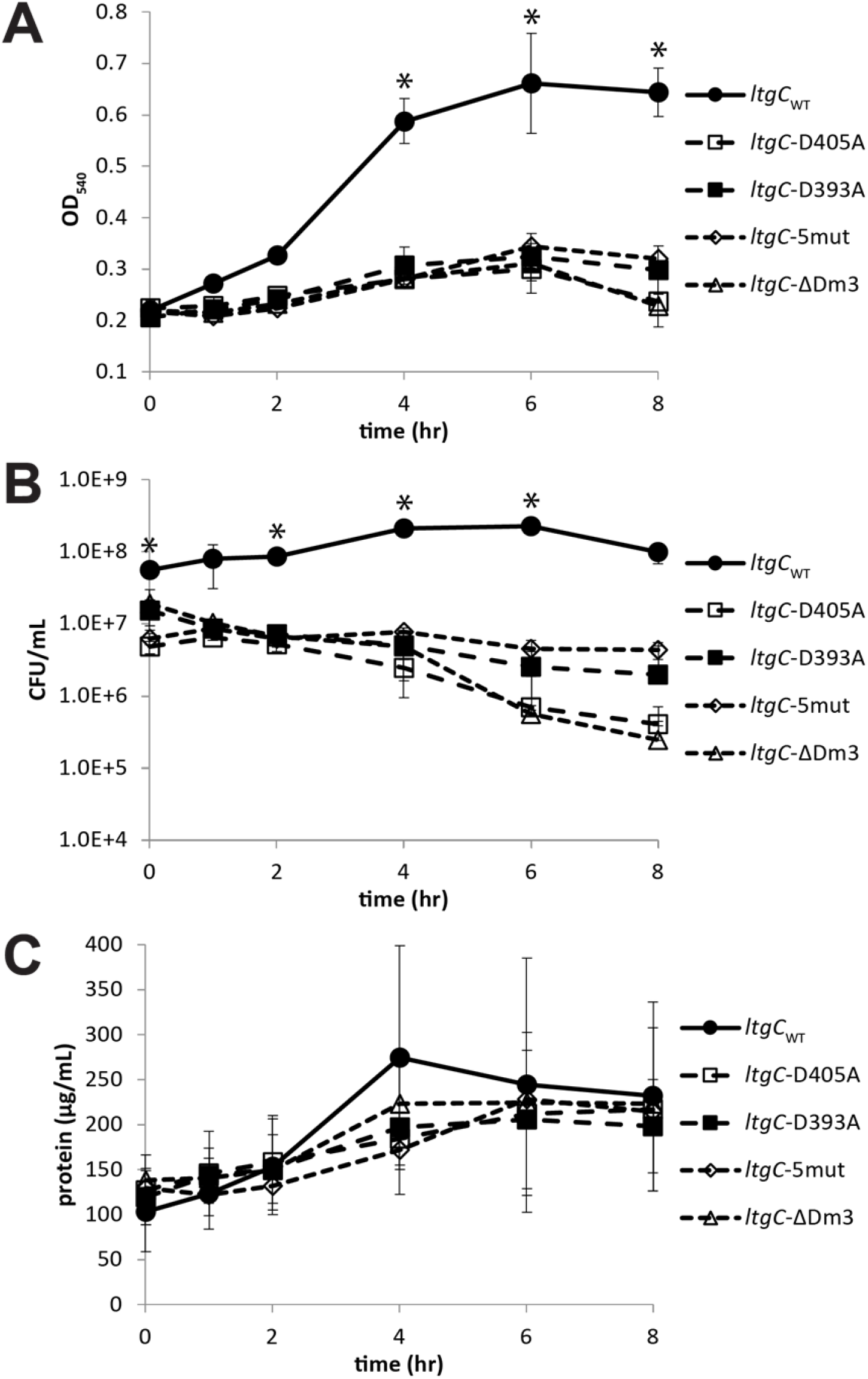
Growth characteristics of *ltgC* mutants. The growth of *ltgC* mutants over 8 hours was measured by optical density at 540nm (a), colony forming units (b), and total protein (c). Optical density and CFU/mL amounts over time are lower for the *ltgC* mutants compared to wild-type due to cell separation defects. There was no significant difference in the amount of protein incorporated into cells, indicating that *ltgC* mutants do not have a growth defect.

### LtgC biochemical function *in vitro*

We expected that the active site mutations would reduce or eliminate enzymatic function of LtgC, but it was unclear whether mutations in domain 3 would have similar effects. Therefore, we purified the proteins and evaluated their activity in zymogram assays and peptidoglycan degradation assays. In addition to using the enzyme from *N. gonorrhoeae* strain FA19, we also cloned and expressed *ltgC* from gonococcal strain MS11, a strain used in multiple analyses of peptidoglycan metabolism [4, 11, 13, 17]. For the zymograms, *Micrococcus luteus* cells were suspended in the polyacrylamide, and the purified LtgC proteins were electrophoresed through the gel. The gel was subjected to multiple rounds of washing in buffer to allow the proteins to refold; then it was stained with methylene blue. Zones of clearing can indicate that the enzyme has degraded the peptidoglycan in the gel, though the method is prone to yielding false positive reactions for some proteins [18, 19]. WT LtgC from FA19 or MS11 showed a substantial zone of clearing. However, each of the mutant proteins showed a markedly fainter zone of clearing (Fig 4), with the catalytic site substitutions showing the least amount of clearing and the 5mut protein somewhat reduced in clearing.

**Figure 4:**
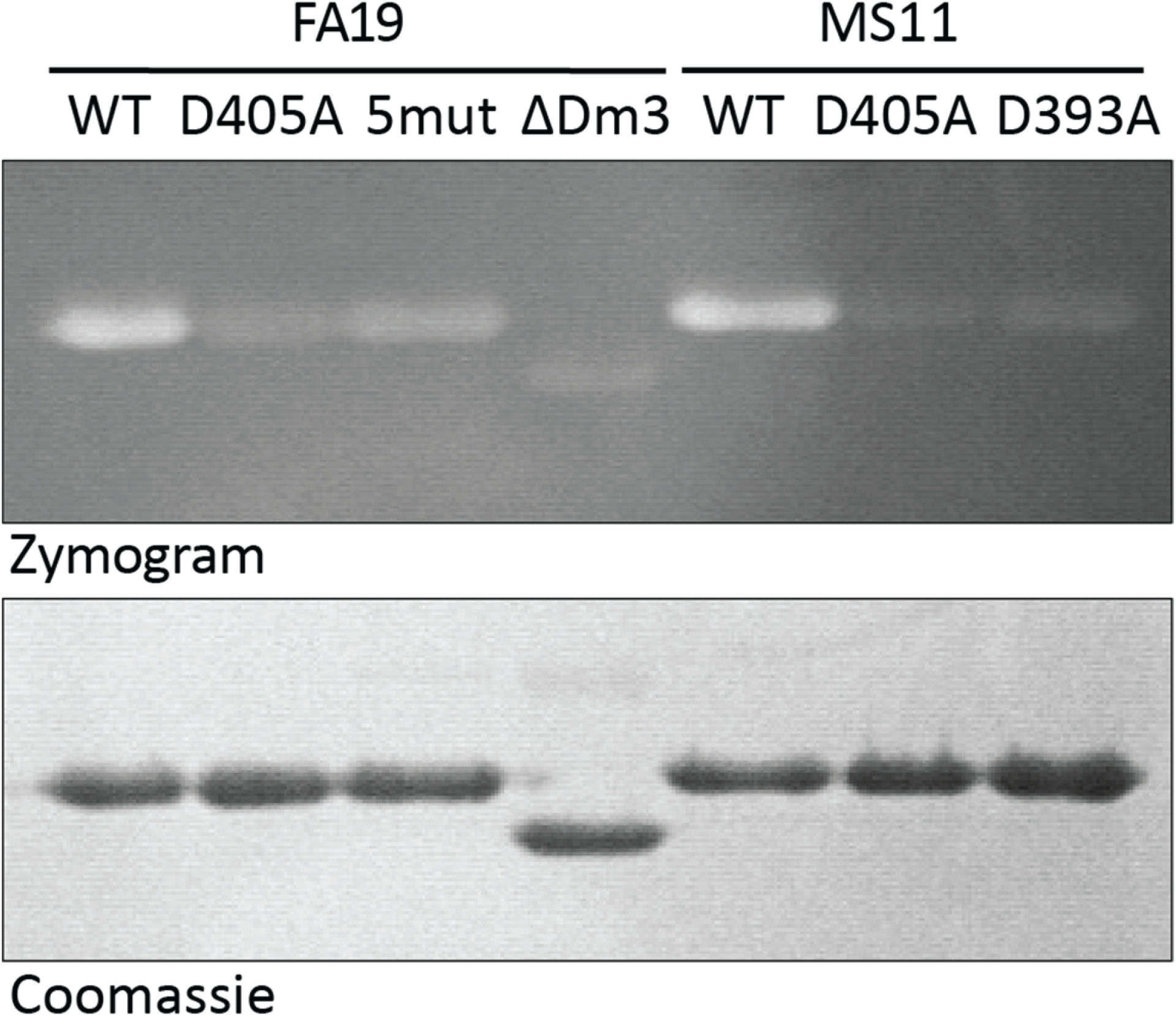
Zymogram analysis of LtgC activity. LtgC proteins with C-terminal His6-tags were purified, and 5µg of nickel affinity purified protein from each strain was electrophoreses on a 10% acrylamide gel containing 0.2% lyophilized *Micrococcus* cells. The gel was renatured overnight and was then stained with methylene blue (top). The gel was later stained with Coomassie to visualize proteins and ensure equal protein amounts (bottom). The zones of clearing show that wild-type LtgC from both FA19 and MS11 degrade peptidoglycan. Some activity can be seen from the LtgC-5mut, and all of the mutants showed less clearing than seen with the wild types.

To measure quantitatively the effects of the mutations on enzymatic function, we tested the abilities of the proteins to degrade ^3^H-labeled gonococcal peptidoglycan*. N. gonorrhoeae* carrying a mutation in *pacA* was metabolically labeled by growth in medium containing 6-[^3^H]-glucosamine and lacking glucose to produce non-acetylated peptidoglycan labeled at both GlcNAc and MurNAc. The peptidoglycan was purified by the boiling SDS method and was used as a substrate for LtgC reactions [20]. The purified LtgC proteins were added to labeled peptidoglycan, and at each time point, the samples were centrifuged, and peptidoglycan solubilization was measured as radioactivity in the supernatant. As expected, the WT enzymes from FA19 or MS11 showed significant activity, and the catalytic site mutants showed minimal solubilization of peptidoglycan (Fig 5). Of particular interest, the domain 3-defective enzymes (5mut and ΔDm3) gave intermediate values, reflecting partial activity and mirroring the results from the zymogram analysis.

**Figure 5:**
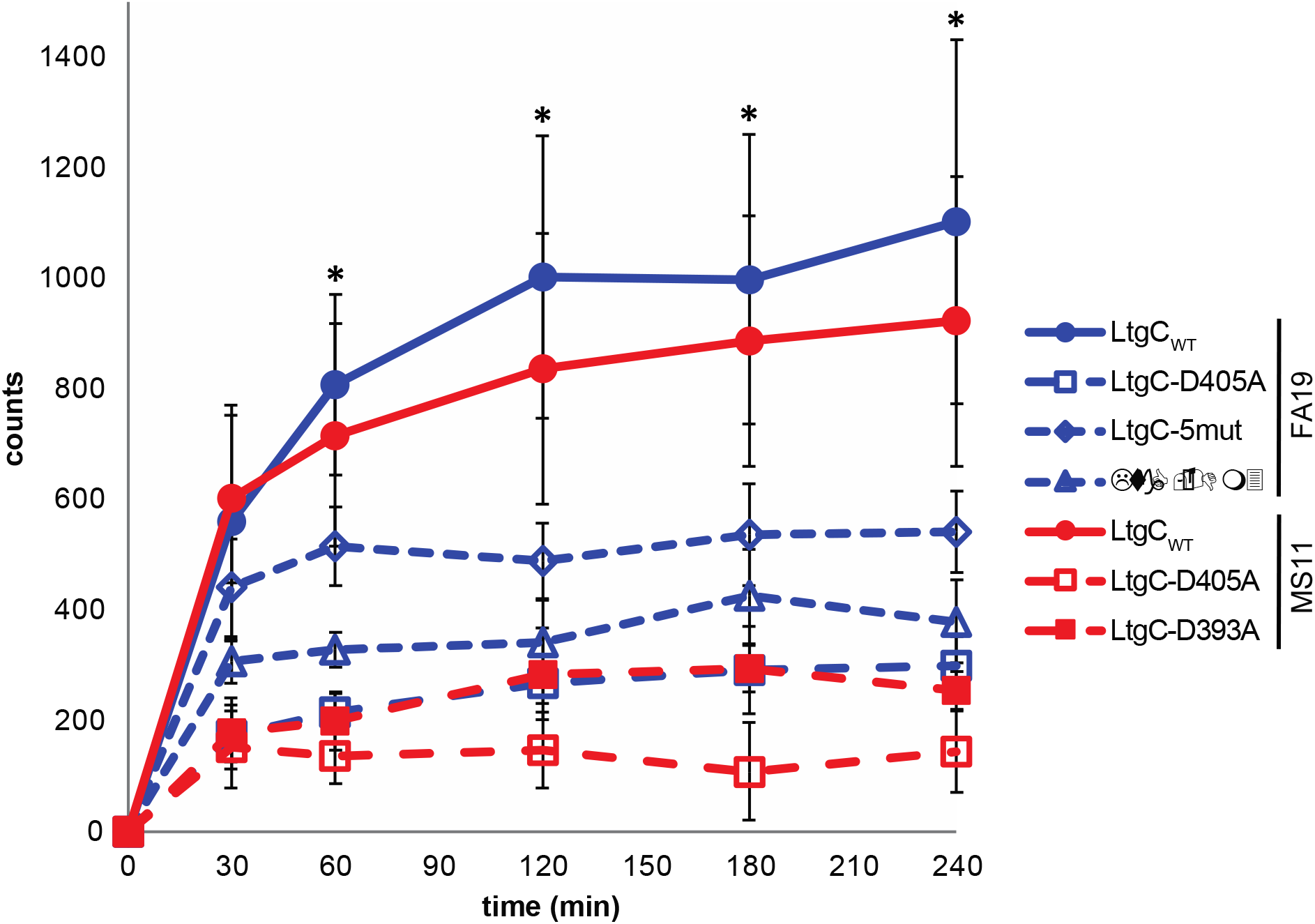
Solubilization of radiolabeled peptidoglycan by LtgC. Radiolabeled MS11 *ΔpacA* peptidoglycan was combined with 5 nmoles of purified LtgC. Samples of the reactions were collected at the indicated times, and the insoluble, macromolecular PG was precipitated by the addition of TCA and collected by centrifugation. The soluble fragments were quantified by scintillation counting. Values are the average of three independent experiments and error bars show standard deviation. Asterisk (*) indicates where LtgC_WT_ was significantly different (*p* < 0.05) than all mutants as determined by Students’ *t*-test. LtgC proteins derived from strain FA19 are shown in blue, and LtgC proteins derived from strain MS11 are shown in red.

### Release of peptidoglycan fragments during growth

*N. gonorrhoeae* releases small peptidoglycan fragments as the bacteria grow, with the most abundant fragments being the peptidoglycan monomers, GlcNAc-anhydro-MurNAc-tripeptide and GlcNAc-anhydro-MurNAc-tetrapeptide [5, 21]. Other abundant fragments released include peptidoglycan dimers, in which two monomers are linked together either through peptide crosslinks or through glycan linkage. Gonococci also release a tetrasaccharide-peptide, which is similar to a glycosidically-linked dimer, but with only one peptide. Free disaccharides of GlcNAc-anhydro-MurNAc are released, and free anhydro-MurNAc monosaccharide is sometimes detected [21]. The WT FA19 and the derivative expressing LtgC-HA showed the expected profile of released peptidoglycan fragments (Fig 6). Mutation of one of the catalytic residues (LtgC-D405A) resulted in a strain that did not release either free disaccharide or tetrasaccharide-peptide (Fig 6A). It did, however, release more large peptidoglycan fragments, typical of strains that show increased cell lysis [22, 23]. This fragment release profile was similar to that of a mutant with a deletion of the 5’ coding region of *ltgC* [9].

**Figure 6:**
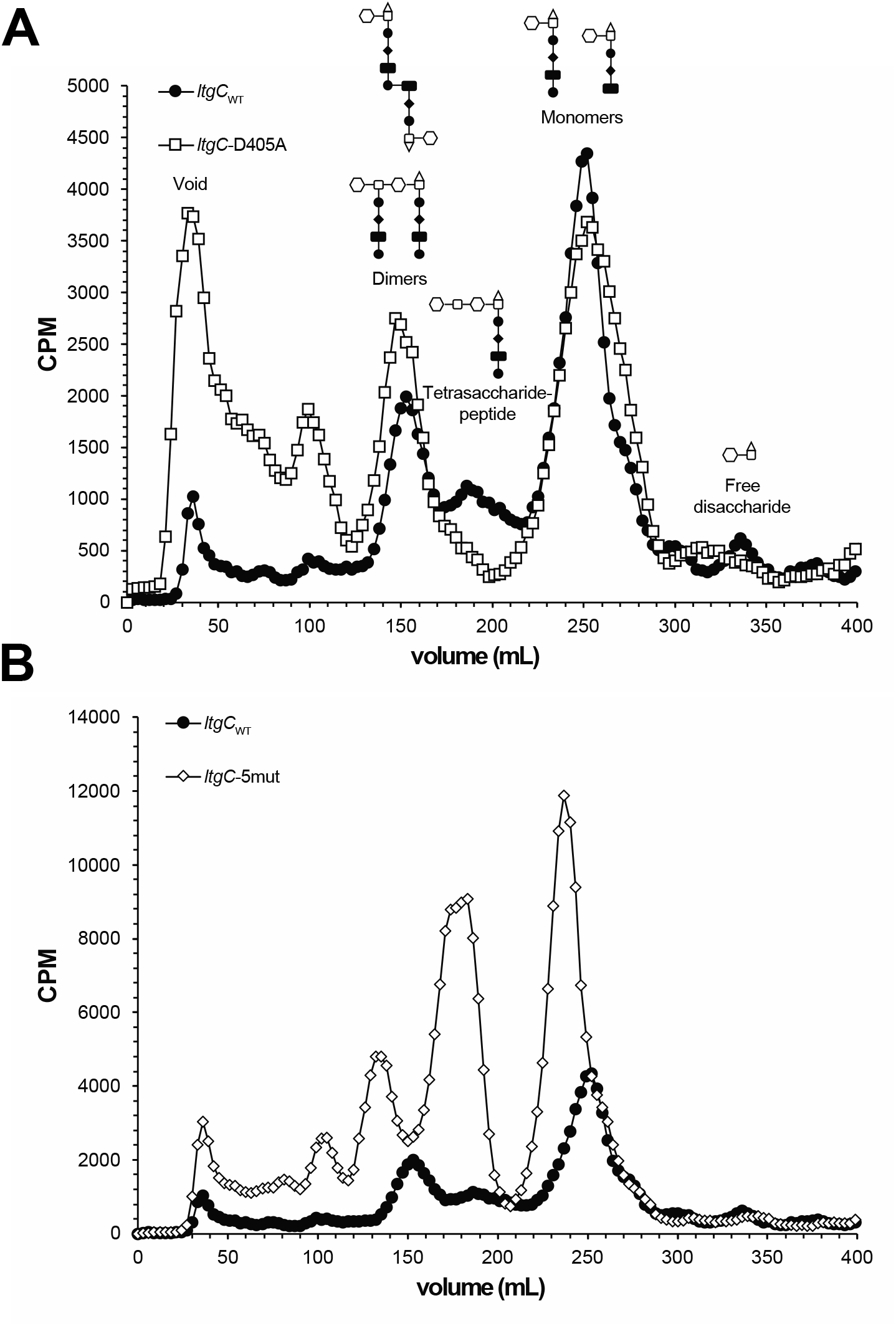
Fragment release profiles of *ltgC* mutants. Strains were pulse labeled with [6-^3^H]-glucosamine, and peptidoglycan fragments released by growing gonococci were separated by size-exclusion chromatography. The *ltgC*-D405A active-site point mutation resulted in loss of the tetrasaccharide peptide and the free disaccharide peak. Mutation of charged surface residues in domain 3 (LtgC-5mut) resulted in an increase of both the size and abundance of peptidoglycan-derived fragments released.

The 5mut strain had a very interesting peptidoglycan fragment release phenotype. Similar to the phenotype of the *ltgC*-D405A mutant, the release of free disaccharides was diminished. However, much larger effects were seen for the other peptidoglycan fragments (Fig 6B). There was an increase in the release of monomers, and they migrated at a size consistent with GlcNAc-anhydro-MurNAc-pentapeptide [24]. Peaks near the sizes of tetrasaccharide-peptide and dimers were also shifted to larger molecular sizes. As with the LtgC-D405A mutant strain, FA19 expressing LtgC-5mut released significant amounts of large peptidoglycan fragments consistent with increased cell lysis (Fig 6). These results are consistent with LtgC degrading peptidoglycan strands that have had their peptides removed except for the last one. The D405A mutant would not be able to degrade these long strands and thus not produce free disaccharides nor the tetrasaccharide-peptide.

### Peptidoglycan composition in the sacculus

While it is expected that LtgC acts to degrade PG strands that are removed during cell separation, it may also degrade strands in the sacculus or alter the activity of other peptidoglycan-degrading enzymes. To determine the effects of LtgC on the sacculus, we purified macromolecular peptidoglycan and determined its composition using UPLC/MS. The various *ltgC* mutants showed significant differences in composition of the peptidoglycan sacculus (Table 1). All mutants had decreased amounts of *O*-acetylated disaccharide tripeptides (Tri) and *O*-acetylated disaccharide tetrapeptides (Tetra). These differences could be due to a decreased amount of peptidoglycan acetylation, or they could be due to increased deacetylase activity. In addition, there was an increase in Tri-Tetra dimers in the mutants. The largest difference was a near 3-fold increase in disaccharide pentapeptides (Penta). This result suggests that newly added peptidoglycan is not being crosslinked to the growing sacculus as efficiently in the *ltgC* mutants as in the WT or that D-alanyl-D-alanine carboxypeptidases such as PBP3 or PBP4 have diminished activity.

**Table 1:** Composition of sacculi of *ltgC* mutants. Insoluble peptidoglycan was purified using boiling SDS and proteases. Sacculi were then digested with mutanolysin and treated with mutanolysin to digest the glycan backbone. LC-MS was used to identify digested peptidoglycan fragments.

| Peak | Identity | *m/z* | Charge | WT | *ItgC*-<br>*HA* | Δ*ItgC* | *ItgC*(D405A)-<br>*HA* | *ItgC*(D393A)-<br>*HA* | *ItgC*-<br>*5mut*-<br>*HA* | *ItgC*ΔDom3-<br>*HA* |
| --- | --- | --- | --- | --- | --- | --- | --- | --- | --- | --- |
| 1 | Tri | 871.32 | 1H+ | 4.56 | 4.67 | 6.04 | 4.19 | 6.22 | 3.89 | 7.29 |
| 2 | Tetra | 942.35 | 1H+ | 21.74 | 25.39 | 25.43 | 24.44 | 24.53 | 20.79 | 24.00 |
| 3 | Tri(OAc) | 913.34 | 1H+ | 9.84 | 10.42 | 8.71 | 7.76 | 6.86 | 7.34 | 7.24 |
| 4 | Penta | 1013.38 | 1H+ | 4.25 | 5.43 | 14.95 | 15.19 | 13.87 | 12.02 | 16.62 |
| 5 | Tetra(OAc) | 984.47 | 1H+ | 20.28 | 22.08 | 15.55 | 15.33 | 18.42 | 17.15 | 15.19 |
| 6 | Tri-Tetra | 897.74 | 2H+ | 1.63 | 0.92 | 2.94 | 2.36 | 2.17 | 3.37 | 2.34 |
| 7 | Tetra-tetra | 933.45 | 2H+ | 4.44 | 4.43 | 6.06 | 5.53 | 7.79 | 3.87 | 4.55 |
| 8 | Tetra-tetra(OAc) | 954.68 | 2H+ | 6.03 | 4.06 | 4.18 | 3.86 | 4.46 | 3.81 | 3.83 |

### STORM localization LtgC in gonococcal cells

We used super-resolution microscopy to examine the location of LtgC in gonococcal cells. A 3xFLAG tag was translationally fused to the 3’ end of *ltgC* in strain MS11 to allow detection of the protein. Fluorescence microscopy using DAPI to stain gonococcal DNA shows that the FLAG tag results in somewhat decreased cell separation, though not to the extent seen in *ltgC* mutants (Fig 7A). LtgC-3xFLAG was localized to the septum between diplococci or to the septa between cells growing in a cluster. The LtgC-D405A mutant showed similar localization with LtgC detected at the septa (Fig 7B).

**Figure 7:**
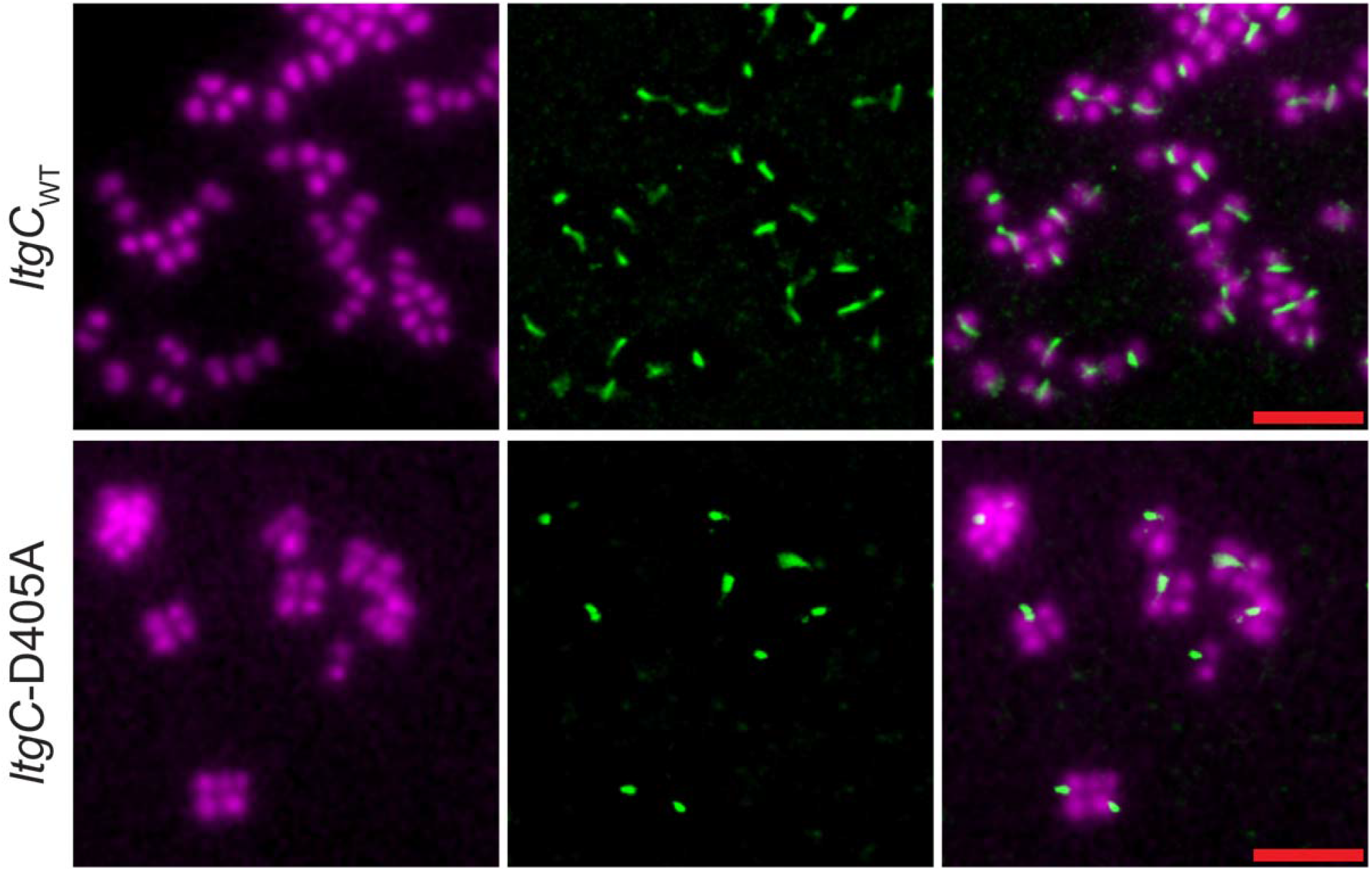
Stochastic optical reconstruction microscopy depicting the localization of LtgC_WT_ and LtgC-D405A active-site point mutant with C-terminal FLAG3-tag fusion (green). DAPI staining showing the location of DNA and thus the gonococcal cells is shown in purple. In the merged micrographs, LtgC can be seen localizing to the septum of dividing cells for both the WT and mutant bacteria. LtgC cellular localization is not dependent on enzymatic activity. Red scale bar = 2µm.

### Survival in human blood

A defect in cell separation in an *N. meningitidis nlpD* mutant was shown to result in a deficiency in survival in a human whole blood infection model [14]. Such a model is also relevant to *N. gonorrhoeae* since gonococci can cause disseminated infection [25], and the strain used here, FA19, was isolated from a patient with a disseminated infection [26, 27]. Therefore, we tested the ability of the *ltgC* mutants to survive in hirudin-treated human blood. Upon inoculation, all the strains declined in CFU numbers (Fig 8). After 2h, the WT strain and the ΔDm3 mutant begin to grow in the blood. By contrast, the mut5 and D405A mutants decreased over 10-fold in 2h and remained near that level throughout the 6h time period. All of the mutants exhibited significantly poorer survival compared to the WT. Interestingly, the ΔDm3 mutant showed an intermediate level of survival (Fig 8).

**Figure 8:**
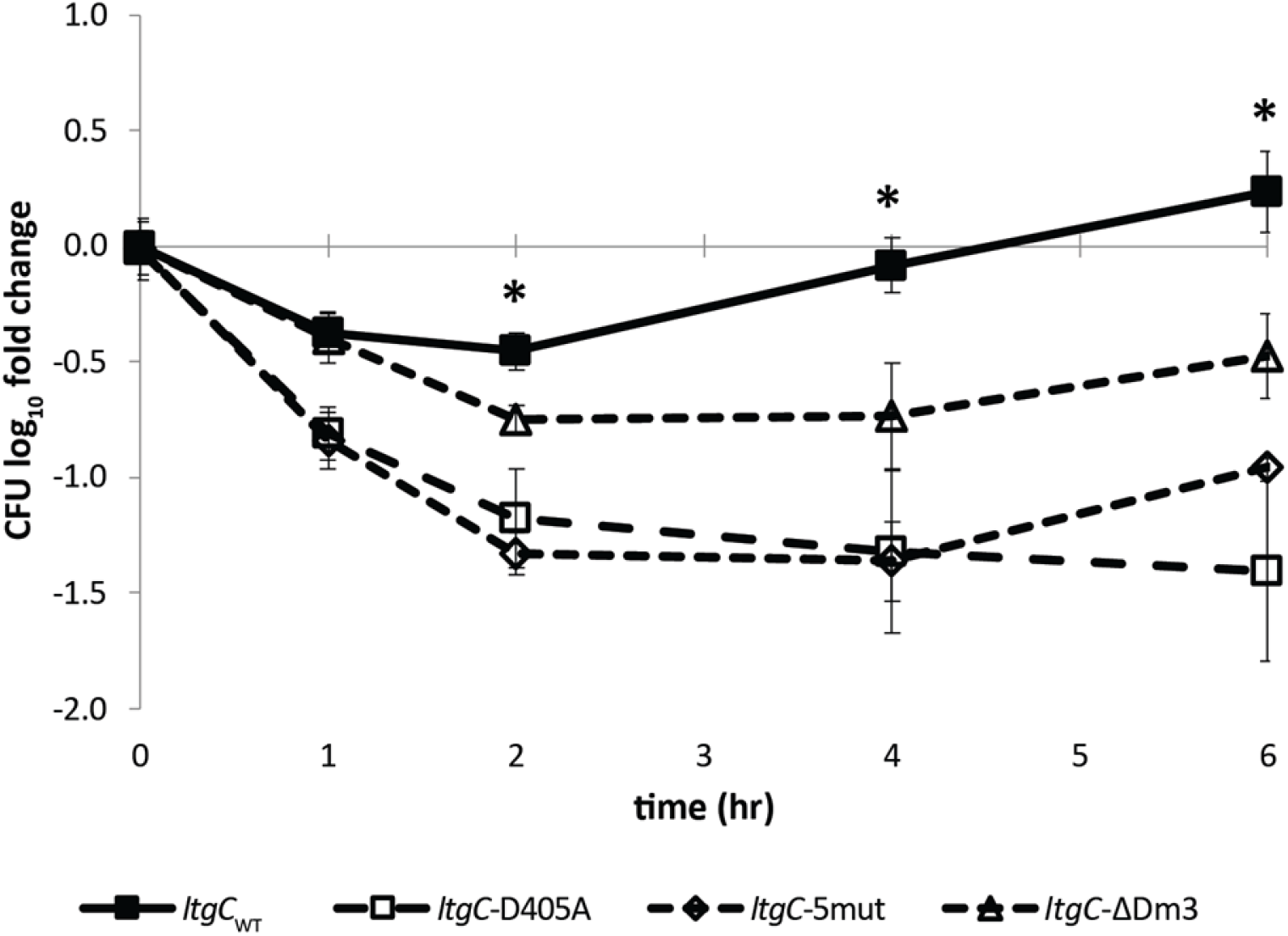
Survival of *ltgC* mutants in human blood. Human whole blood with hirudin was inoculated with an equivalent of OD_540_=0.2 of bacteria, and CFUs/mL were determined over 6 hours. Due to differences in cell separation, CFU counts do not represent the number of viable bacteria. Therefore, bacterial survival in blood is reported as the log_10_ (CFUs / CFUs at inoculation). Error bars indicate standard deviation. Asterisk (*) indicates that *ltgC*_WT_ was significantly different than all mutants (*p*-value < 0.05).

### LtgC interacts with AmiC in BACTH assays

The *N*-acetylmuramyl-L-alanine amidase AmiC is required for cell separation in *N. gonorrhoeae*, like LtgC [11]. We used the bacterial two-hybrid (BACTH) assay to determine if LtgC interacts with AmiC and if domain 3 is important for the interaction. Indeed, LtgC fused to a subunit of adenylate cyclase interacted with AmiC fused to the other subunit of adenylate cyclase resulting in enzymatic activity that was detected in *E. coli* transformants containing the fusions (Fig 9A). The fusions also interacted when the adenylate cyclase subunits they were attached to were swapped, suggesting that the interaction was not an artifact of the particular construct used. Fusions to the *mut5* and Δ*Dm3* mutant alleles of *ltgC* were also used. These mutant versions of the protein also showed interaction with AmiC. However, a quantitative assay using the LacZ reporter showed reduced interaction, with the *mut5* allele showing a moderate reduction and the Δ*Dm3* allele showing a much larger reduction (Fig 9B). These data indicate that domain 3 is important for AmiC-LtgC interaction but that other portions of LtgC also interact with AmiC.

**Figure 9:**
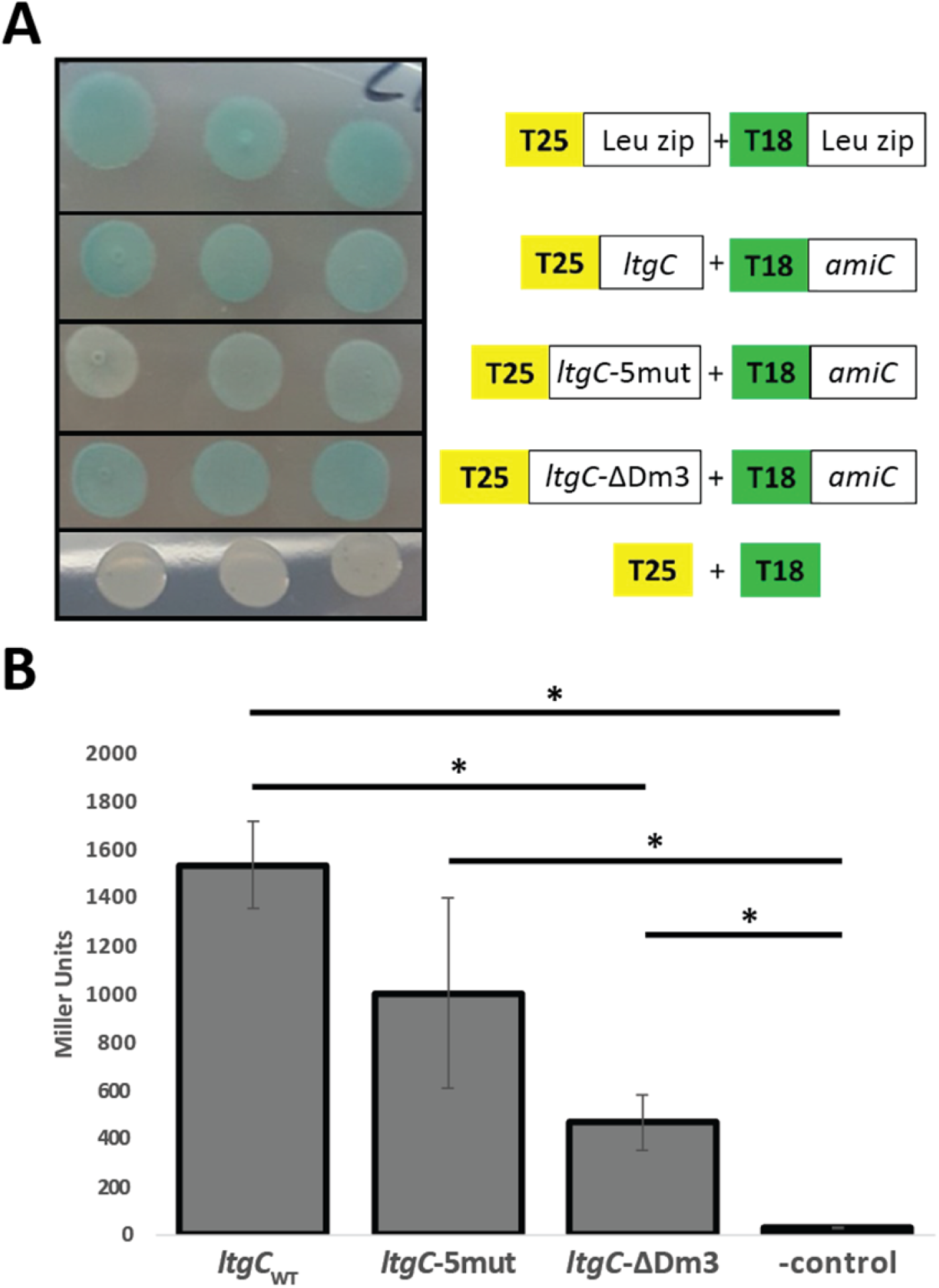
Domain 3 of LtgC is involved in interaction with AmiC. (A) Results from colorimetric assay using the BACTH system to test for interaction between AmiC and LtgC domain 3 mutants. *E. coli* BTH101 cells were co-transformed with the respective plasmids and spotted on LB X-Gal plates. Images were taken after incubation at 30°C for 2 days. Positive control (top row), LtgC-T25+AmiC-T18 (2^nd^ row), LtgC5mut-T25+AmiC-T18 (3^rd^ row), LtgCΔdom3-T25+AmiC-T18 (4^th^ row) and negative control (bottom row). Blue color indicates that interaction occurs between the two proteins and white color indicates no interaction. (B) Results from β-galactosidase assays with the same strains used for the colorimetric assay. Increase in β-gal activity relative to the negative control indicates protein interaction. Error bars depict standard error. Asterisk (*) indicated *p* < 0.05.

## Discussion

Mutation of *ltgC* in *N. gonorrhoeae* or *N. meningitidis* was shown previously to result in loss of normal cell separation, causing the bacteria to grow in clusters [9, 10]. The present study demonstrates that loss of domain 3 of LtgC or changes in the surface residues of domain 3 of LtgC also causes diminished cell separation, even though those mutants (ΔDm3 and 5mut) retain significant enzymatic activity (Fig 2I-2L, Fig 5). It is unclear exactly why lytic transglycosylase activity is needed for cell separation. Some studies suggest peptidoglycan strands directly connect the daughter cells even after endopeptidase and amidase processing and those strands must be degraded. Other studies suggest periplasmic crowding and osmotic pressure resulting from removed but undegraded glycan strands result in cell separation defects [28, 29]. LtgC is thought to degrade glycan strands following or in concert with amidase action to remove peptides from peptidoglycan. The peptidoglycan release data in Fig 6, data from Cloud and Dillard [9] showing a requirement for LtgC for producing free disaccharides and tetrasaccharide peptide, and data from Lenz et al. [30] demonstrating the necessity for AmiC for producing tetrasaccharide peptide, are all consistent with this idea.

In other bacterial species where LtgC homologs (often referred to as MltA) have been characterized, the enzyme has been found to form protein-protein interactions with a scaffolding protein, MipA [31]. Neither *N. gonorrhoeae* nor *N. meningitidis* has a MipA homolog. However, gonococcal LtgC does exhibit protein-protein interactions, interacting with the amidase, AmiC (Fig 9). AmiC is the other peptidoglycan-degrading enzyme required for gonococcal or meningococcal cell separation, along with its activator NlpD [11, 12, 14]. We show here that AmiC binds to LtgC in bacterial 2-hybrid assays and that mutations in domain 3 of LtgC lead to significantly reduced interactions (Fig 9). These results suggest that AmiC binds to LtgC at domain 3 and at a distal site. Alternatively, it is possible that the domain 3 mutations alter a different region of LtgC involved in AmiC binding. The other characterized LtgC homolog that has a domain 3 is from *Acinetobacter baumannii* [32]. In crystal structures, domain 3 was found to function in protein dimerization. In contrast to *N. gonorrhoeae*, *A. baumannii* has a MipA homolog (GenBank: SSM88690.1) and may use MipA to facilitate interaction with amidases or other cell separation proteins.

Consistent with its role in cell separation, LtgC is found at the septum between diplococci or clustered cocci (Fig 6). This localization is similar to that of the amidase activator NlpD in *N. meningitidis* [14]. The LtgC localization is somewhat different from that of other lytic transglycosylases. LtgA shows different localization at different points in growth, spread around gonococcal cells when they are monococci but found at the septum during division [20]. Lytic transglycosylase LtgD is found around the cell at all points in growth [20]. Amidase AmiC shows a localization pattern similar to LtgA, present at the septum only in dividing cells [14].

All of the *ltgC* mutants showed a defect in survival in human blood. This phenotype is similar to that observed with an *nlpD* mutant of *N. meningitidis* that was similarly deficient in cell separation [14]. The reasons for the reduced survival are not known, but defects with the outer membrane seem likely. Since cell separation is reduced, the outer membrane must surround the large cluster of cells. Studies of a gonococcal *amiC* mutant defective in cell separation demonstrated increased sensitivity to deoxycholate, suggesting an outer membrane defect [11]. Such a defect could make the bacteria more sensitive to various innate immune components.

Overall, our results suggest a model in which LtgC functions along with AmiC to degrade peptidoglycan strands at the septum during cell separation (Fig. 10). First, an endopeptidase, such as PBP3 (DacB) or PBP4 (PbpG) cuts peptide crosslinks. AmiC is part of a protein complex, including its activator NlpD and the lytic transglycosylase LtgC. LtgC directly binds to AmiC, and the binding is partly facilitated by domain 3 of LtgC. AmiC removes the peptides from MurNAc residues, except for the one on the anhydro-MurNAc at the end of the strand [30]. This process generates free tripeptides and tetrapeptides, and a portion of these peptides are released from the bacteria (Fig. 10A). LtgC degrades the denuded glycan strand into anhydro-MurNAc disaccharides and a single tetrasaccharide peptide from the end of the strand. Quantities of these fragments are also released by the bacteria (Fig. 6A). Without LtgC, AmiC is still able to release free peptides (Fig. S1), but the glycans remain uncut and retain a peptide at anhydro-MurNAc, such that neither free disaccharide nor tetrapeptide-peptide is released (Fig 6A, Fig. 10B). A loss of LtgC activity or LtgC binding to AmiC results in the bacteria remaining in a cluster sharing one cell wall and one outer membrane (Fig 2) and becoming highly susceptible to killing by the host innate immune system.

**Figure 10:**
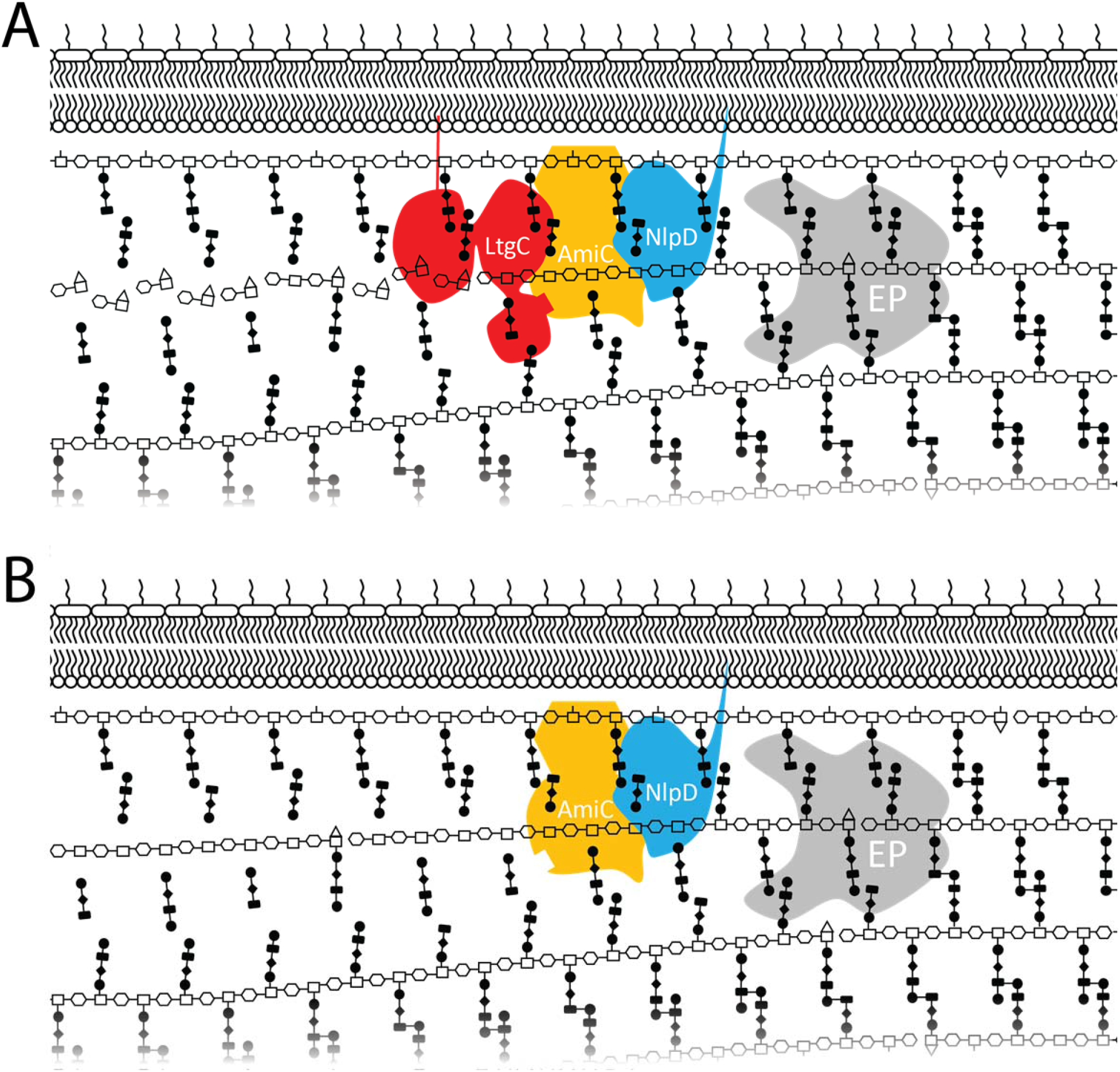
Model of peptidoglycan breakdown at the septum during cell separation. (A) Endopeptidase (EP) activity from PBP3 or PBP4 is required for cutting peptide crosslinks before the amidase AmiC can remove peptides from the cell wall. AmiC is in a complex with its activator NlpD and with lytic transglycosylase LtgC. AmiC removes all of the peptides except the one on the anhydro-MurNAc at the end of the strand. After AmiC removes the peptides, LtgC degrades the naked strand into GlcNAc-anhydro-MurNAc disaccharides and a tetrasaccharide peptide that comes from the end of the strand. Each of these fragments is released into the extracellular milieu. (B) Without LtgC activity, the glycan strands remain long and disrupt cell separation processes, possibly by affecting osmolarity of the periplasm or remaining attached to cell wall in both daughter cells.

## Methods

### Bacterial Growth Conditions and Strains

*Neisseria gonorrhoeae* strains were derived from either strain FA19 or MS11. Cells were grown on GCB agar plates with Kellogg’s supplements in the presence of 5% CO_2_ or in complemented gonococcal base liquid (GCBL) with 0.042% sodium bicarbonate and Kellogg’s supplements. *Escherichia coli* strains were grown in lysogeny broth (LB) [1% tryptone, 0.5% yeast extract, 0.5% NaCl]. FA19 *ltgC-HA* strains were created by making constructs containing an in-frame HA-tag fusions and a downstream kanamycin cassette. The ΔDom3 construct was made by deleting the sequence between codon 166 and 228 and adding an RGR with a SacII restriction site. The *ltgC-5mut* strain contains the mutations K179A, L181A, R183A, L227A, and P228A. Point mutations were made by overlap extension. The resulting plasmids were then transformed into *N. gonorrhoeae* FA19 and selected on kanamycin plates.

To create recombinant LtgC N-terminal hexahistidine fusion proteins and corresponding mutants from FA19 and WT LtgC from MS11, oligos LtgC-NS-NdeI-F (5’-T<u>TC ATA TG</u>A GCA GGA GCA TCC AAA CCT TTC CG) and LtgC-XhoI-R (5’-T<u>TC TCG AG</u>T CAC GGG CGG TAT TCG GGC) were amplified from the above strains and ligated into pET28b at the NdeI/XhoI restriction sites. The D405A mutation in MS11 LtgC was introduced by overlap extension by amplifying MS11 DNA with both LtgC-Kpn-F (5’-TT<u>G GTA CC</u>T TAT GAA AAA ACA CCT GCT CCG) and LtgC-D405A-R (5’-AAA ATA AGC CAC GCG CAC CGC G), and LtgC-D405A-F (5’-GTG CGC GTG GCT TAT TTT TGG GGT TAC G) and LtgC-Sac-R (5’-TTG AGC TCT CAC GGG CGG TAT TCG GGC). The resulting PCR products were then used as a template with primers LtgC-Kpn-F and LtgC-Sac-R and ligated into pPK33 [33]. Similarly, the MS11 D393A mutation was introduced using the product of oligos LtgC-Kpn-F and LtgC-D393A-R (5’-CGC TGC CTG TAG CCT GCG CCA TAA), and LtgC-D393A-F (5’-TTA TGG CGC AGG CTA CAG GCA GCG) and LtgC-Sac-R as a template with LtgC-Kpn-F and LtgC-Sac-R and the resulting product was ligated into pPK33 at the KpnI/SacI sites. The MS11 derived LtgC point mutant constructs were then amplified with LtgC-NS-NdeI-F and LtgC-XhoI-R and inserted into pET28b at the NdeI/XhoI sites. The resulting pET28b plasmids were transformed into *E. coli* BL21 (DE3) and selected with kanamycin.

To create constructs with a C-terminal 3XFLAG sequence at the native site, *ltgC* from wild-type and mutants was amplified with LtgC-SacI-Start-F (5’-T<u>TG AGC TC</u>A TGA AAA AAC ACC TGC TCC GCT CC) and LtgC-EcoRI-R (5’-A<u>GA ATT C</u>CG GGC GGT ATT CGG GCT TCA TGC) and ligated into the 3XFLAG containing vector pMR100 (Ramsey ME, *et al.* 2014) at the SacI/EcoRI sites. The downstream homology region was amplified using primers LtgC-DS-HindIII-F (5’-TTT <u>AAG CTT</u> GCA AAA CAA TGC CGT CCG AAG C) and LtgC-DS-Xho-R (5’-T<u>TC TCG AG</u>G CTT TTG TCT GAA AAA GAC GGC GC) and inserted at the HindIII/XhoI sites.

To create plasmids for the bacterial two hybrid assay, primers were designed to amplify the genes of interest without their signal sequence for insertion into plasmids pKT25 and pUT18C [34]. The pUT18C-*amiC* plasmid was made by amplifying *amiC* with amiC F XbaI (5’-AAA <u>TCT AGA</u> GAA AAC GGT ACG CGC CCC GCA G) and amiC R KpnI (5’-TTT <u>GGT ACC</u> GGG CGG AAG TAG GTT TAT GC) and inserting the digested fragment into pUT18C at the XbaI/KpnI sites. The pKT25 with *ltgC* and *ltgC* mutants were amplified with ltgC F XbaI no sig (5’-GCG <u>TCT AGA</u> GCA AAG CAG GAG CAT CCA AAC) and ltgC R XmaI no sig (5’-ATT CCC <u>GGG TCA</u> CGG GCG GTA TTC GGG CTT C) and inserted into pKH25 at the XbaI/XmaI sites.

### Thin-section Electron Microscopy

*N. gonorrhoeae* strains were grown on GCB agar overnight and used to seed 3 mL GCBL cultures at OD_540_=0.2. Cultures were grown for 3 hours with aeration to achieve mid-log phase. Cells were harvested by centrifugation and washed with phosphate-buffered saline (PBS) before suspension in a fixative solution of 2% paraformaldehyde and 2.5% glutaraldehyde in a 0.1 M phosphate buffer. Sample preparation and imaging was performed by University of Wisconsin-Madison Medical School Electron Microscope Facility.

### Quantification of Cell Growth

Gonococcal strains were grown on GCB agar plates overnight and diluted to OD_540_=0.2 and grown with aeration at 37°C. At each timepoint, a portion of culture the culture was used to determine OD_540_, CFUs were determined by serial dilution, and protein concentration was determined by Bradford Assay.

### Purification and *In Vitro* Activity of Recombinant Proteins

#### Protein Purification

Overnight cultures were used to seed 1 L culture 1:200. When cultures reached OD_540_=∼0.7 they were transferred to 30°C and induced with 1.0 mM IPTG for 3 hours, washed with 1XPBS, and stored at −80°C. Cells were suspended in nickel buffer (NB) [300 mM NaCl, 20 mM Tris-HCl pH 8] with 20 mM imidazole and lysed twice by French press at 12,000 PSI. Insoluble cell debris was removed by centrifugation an 18,000 x *g* and the cleared lysate was incubated with 600 µL of Ni-NTA slurry at 4°C. On a Ni-NTA resin was transferred to a chromatography column, washed with 10 mL of NB with 20 mM imidazole and eluted with successive 1.2 mL elutions of NB with 60 mM, 100 mM, 250 mM, and 250 mM of imidazole. Elutions were combined and dialyzed against NW with 50% imidazole and stored at −80°C. *Zymogram Analysis*: A 10% acrylamide SDS-PAGE gel was made containing 0.2% (w/v) lyophilized *Micrococcus luteus* cells to act as a substrate. Purified proteins (5 µg) were run at 75 volts at 4°C. To renature proteins, the gel was incubated with a renaturing solution [25 mM Tris-HCl pH=8, 0.5% Triton X-100] overnight at RT with agitation. Fresh renaturing solution was then added, and gel was incubated at 37°C for 30 min. To enhance the visibility of the zones of clearing, the gel was stained for 1 hour with 0.1% methylene blue dissolved in 0.01% (w/v) KOH and destained with water before imaging. The gel was then stained with Coomassie and destained before imaging to visualize proteins.

#### Digestion of Radiolabeled Peptidoglycan

Radiolabeled peptidoglycan was made from a *pacA* deletion mutant lacking *O*-acetylation (KH530) as described by [35, 36]. Briefly, 10 µCi/mL D-[6-^3^H]-glucosamine was added to growing gonococcal cultures seeded at OD_540_=0.25 with a pyruvate carbon source for 45 min. Non-peptidoglycan contamination was removed using the boiling-SDS method and pronase treatment. Non-radiolabeled peptidoglycan was purified using the boiling-SDS-method and pronase treatment. To assess the ability of purified proteins to digest insoluble peptidoglycan sacculi, 700,000 CPM of radiolabeled peptidoglycan was added to 0.2µg of purified protein in a 1.1 mL reaction in a 25 mM Tris-HCL pH=7.5 buffer and incubated at 37°C. At each time point, 200 µL of each reaction was mixed with 500 µL of 20% (v/v) trichloroacetic acid (TCA) and 100 µg (20 µL) of non-radiolabeled peptidoglycan to act as a carrier. Reaction mixtures were placed on ice for 30 minutes and then centrifuged at 45,000 x *g* for 30 min. A 500 µL aliquot of the supernatant containing digested peptidoglycan was used for liquid scintillation counting.

### Peptidoglycan Fragment Release

Metabolic labeling of peptidoglycan in *N. gonorrhoeae* and quantitative analysis of released fragments was performed as previously described [37].

### Imaging by Stochastic Optical Reconstruction Microscopy

Imaging and analysis performed as described previously [20].

### Bacterial Survival in Human Whole Blood

Blood was obtained from volunteers under a protocol approved by the University of Wisconsin-Madison Health Sciences Institutional Review Board (Protocol 2017-1179). Blood was collected in vacutainer tubes containing hirudin, to prevent clotting but not interfere with complement activity. Bacteria grown overnight on GCB plates was suspended in GCBL and quantified by optical density at 540 nm. A pellet of bacteria equivalent to seed a GCBL culture at OD_540_=0.2 was mixed with human whole blood at t=0. At each timepoint, 20 µL of infected blood was serial diluted and plated on GCB plates to determine CFUs.

### Protein Interactions by Bacterial Two-hybrid

E. coli BTH101 competent cells were co-transformed plasmids containing T18 tagged and T25 tagged fusion proteins. Transformants were plated on LB agar plates supplemented with ampicillin 100μg/mL and kanamycin 40μg/mL and incubated for 2 days at 30°C. Liquid cultures were started in triplicates for each transformation in LB supplemented with kanamycin 40μg/mL, ampicillin 100μg/mL and 0.5mM IPTG, and incubated overnight at 30°C with rotation. The next day, 2μL of each liquid culture was spotted on LB-Xgal plates supplemented with kanamycin 40μg/mL, ampicillin 100μg/mL and 0.5mM IPTG. After the spots dried, plates were incubated for 2 days at 30°C, checking each day for a change in color. Spots from interacting proteins turn blue and spots from non-interacting proteins are white. pKT25 and pUT18C empty vectors were used as a negative control, and plasmids pKT25-zip and pUT18C-zip (fragments fused to leucine zipper, provided by the kit) served as a positive control.

For quantitative assays, ß-galactosidase assays were conducted adapting the protocol previously described [38]. Briefly, the same overnight cultures used for the colorimetric assay were used for the quantitative assay. 50 µL of overnight cultures were transferred into a flat bottom microtiter plate already filled with 150 µL LB and OD_600_ was measured. In parallel, 200 µL of each culture were transferred into a microcentrifuge tube containing 800 µL of Z buffer (prepared following the recipe detailed in paper). One drop of SDS 0.01% and two drops of chloroform were added and samples mixed thoroughly for 10s to permeabilize cells. Once the chloroform settled at the bottom of the tube, 50 µL of each sample were transferred into a 96-well microplate containing 150 µL of Z buffer. 40 µL of O-nitrophenyl-ß-D-galactosidase (ONPG, 4mg/mL) were added and the enzymatic reaction carried out at 28°C for 15min measuring OD_420_ every 2min in a microplate reader.

## Supplemental Data

**Fig. S1:**
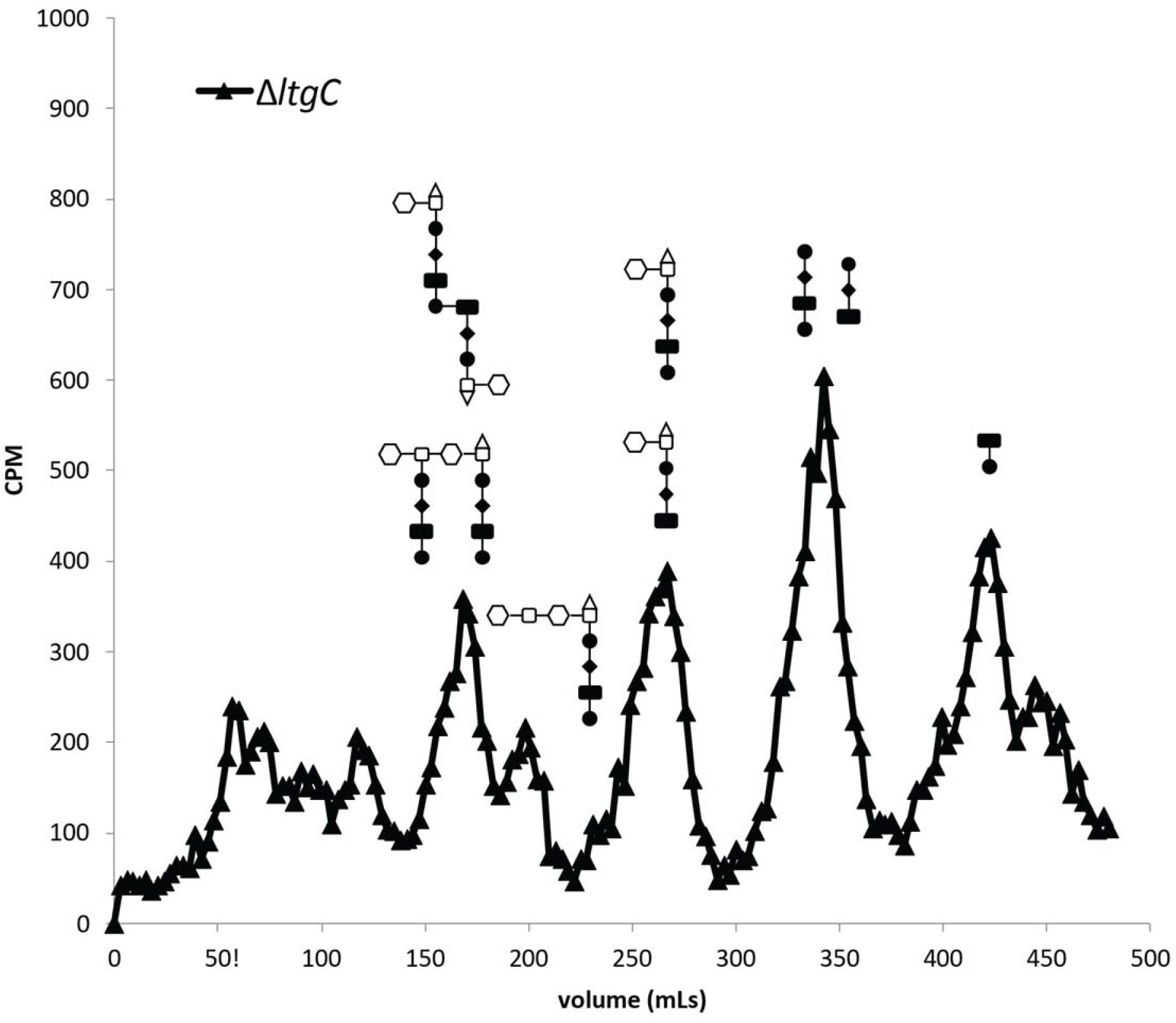
Fragment release profile of Δ*ltgC* mutant KC118. The bacteria were pulse labeled with [2,6-^3^H]-diaminopimelic acid, and peptidoglycan fragments released by growing gonococci were separated by size-exclusion chromatography. Mutation of *ltgC* does not disrupt the release of DAP-containing peptidoglycan fragments, including free peptides generated by AmiC.

